# An efficient gene excision system in maize

**DOI:** 10.1101/2020.05.26.116996

**Authors:** Ning Wang, Maren Arling, George Hoerster, Larisa Ryan, Emily Wu, Keith Lowe, Bill Gordon-Kamm, Todd J. Jones, Nicholas Doane Chilcoat, Ajith Anand

**Affiliations:** Corteva Agriscience, Johnston, IA USA

**Keywords:** *Agrobacterium*, developmentally-regulated promoters, heat-shock promoters, morphogenic genes, marker-free events, rapid maize transformation

## Abstract

Use of the morphogenic genes *Baby Boom* (*Bbm*) and *Wuschel2* (*Wus2*), along with new ternary constructs, has increased the genotype range and the type of explants that can be used for maize transformation. In addition, altering the ectopic expression pattern for *Bbm/Wus2* has resulted in rapid maize transformation methods that are faster and applicable to a broader range of inbreds. However, expression of *Bbm/Wus2* can compromise the quality of regenerated plants, leading to sterility. We reasoned excising morphogenic genes after transformation but before regeneration would increase production of fertile T0 plants. We developed a method that uses an inducible site-specific recombinase (*Cre*) to excise morphogenic genes. The use of developmentally regulated promoters, such as *Ole, Glb1, End2* and *Ltp2*, to drive *Cre* enabled excision of morphogenic genes in early embryo development and produced excised events at a rate of 25%-100%. A different strategy utilizing an excision-activated selectable marker produced excised events at a rate of 53.3%-68.4%; however, the transformation frequency was lower (12.9%-49.9%). The use of inducible heat shock promoters (e.g. *Hsp17.7, Hsp26*) to express Cre, along with improvements in tissue culture conditions and construct design, resulted in high frequencies of T0 transformation (29%-69%), excision (50%-97%), usable quality events (3.6%-14%), and few escapes (non-transgenic; 14%-17%) in three elite maize inbreds. Transgenic events produced by this method are free of morphogenic and marker genes.

## INTRODUCTION

The use of the morphogenic genes *Bbm* and *Wus2* has considerably increased transformation frequencies and reduced genotype dependence in many cereal crops (Lowe et al., 2016;Mookkan et al., 2017;Anand et al., 2018;Lowe et al., 2018). This enabled the development of a rapid transformation method involving direct formation of somatic embryos and T0 plants from immature scutella (Lowe et al., 2018). This approach has facilitated transformation (Lowe et., 2016; Mookkan et al.,2017) and CRISPR/Cas-mediated editing (Chilcoat et al., 2017) in numerous elite maize inbreds, and enabled use of alternate explants, such as embryo slices from mature seeds or leaf segments, for successful maize transformation (Lowe et al., 2016;Lowe et al., 2018). However, ectopic expression of the morphogenic genes often resulted in pleiotropic effects including abnormal shoots/roots and infertile plants (Lowe et al., 2016). The use of promoters that drive high expression levels during the transformation process, but lower expression levels in the vegetative plant, provides one option to ameliorate these problems (Lowe etal.,2018) but the presence of morphogenic genes can still result in some negative effects and is undesirable in commercial products. While fertile T0 plants can be recovered under these conditions, non-visible pleiotropic effects remain a distinct possibility. Similarly, transgenic plants regenerated through *de novo* meristem induction stimulated by morphogenic gene expression also resulted in developmental abnormalities (Maher et al., 2020), and without removal also raise concerns that non-visible pleiotropic effects are possible. Therefore, excising the morphogenic genes is desirable for regenerating healthy plants, for transgene testing and commercial product development. Previously a method using a non-integrating *Wus2* gene expression approach recovered fertile T0 plants free-off morphogenic genes, however this method needed a plant selectable marker gene (SMG) for regenerating events (Hoerster et al., 2020). Here we report an approach that allows excision of both the morphogenic gene and the SMG used in transformation at the same time. As an added benefit this method eliminates any adverse effect from the non-trait genes in commercial products.

Different strategies have been developed for the removal of helper genes following plant transformation, often focused on removing plant selectable markers. One approach is co-transformation with two constructs, one with the SMG and one with the gene of interest. In a transgenic plant with independent insertions of each of these constructs, the selectable marker can be segregated genetically (Hare and Chua, 2002;Puchta, 2003;Darbani et al., 2007;Ling et al., 2016). Alternatively, SMGs can be removed by excision via homologous recombination (Puchta, 2000;Zubko et al., 2000), elimination by transposition (Maeser and Kahmann, 1991;Gao et al., 2015) or, by the use of recombinases to excise unwanted DNA. Several recombination systems have been used to excise SMGs, including *Cre/lox* from bacteriophage P1 (Hoess et al., 1982;Hoess and Abremski, 1985), *Flp/frt* from *Saccharomyces cerevisiae* (Cox, 1983;Senecoff et al., 1985), R/RS from *Zygosaccharomyces rouxii* (Araki et al., 1985), and *Gin/gix* from bacteriophage (Klippel et al., 1988). Recombinases have been delivered via retransformation (Odell et al., 1990;Dale and Ow, 1991), sexual crosses (Bayley et al., 1992;Kilby et al., 1995;Kerbach et al., 2005), or transient expression (Gleave et al., 1999;Kopertekh et al., 2004;Kopertekh and Schiemann, 2005;Jia et al., 2006). In most of these systems excision takes place after the T0 generation and requires screening multiple plants to find one that has undergone successful excision. A design where the SMG and the recombinase genes are on the same construct between the recombination sites has been referred to as “auto-excision” (Verweire et al., 2007;Moravčíková et al., 2008), and allows generation of SMG-free events. By placing the recombinase under the regulation of an inducible/chemical promoter, an expression system that allowed spatial and temporal control (regulated by external or intrinsic signals) was shown to be faster and less resource-intensive (Chong-Pérez and Angenon, 2013;Yau and Stewart, 2013).

We have evaluated three different strategies for auto-excision prior to regeneration to recover stable T0 plants free of morphogenic genes and in some cases the SMG as well: 1) an auto-excision system involving developmentally regulated promoters, 2) an excision-activated marker gene system, and 3) an inducible promoter approach for excising both the morphogenic genes and the SMG. The excision strategies were evaluated to meet key production transformation criteria of 1) high transformation frequency, 2) high quality event (QE, single-copy of T-DNA, backbone and morphogenic gene free) frequency, 3) ability to generate marker-free T0 plants, and 4) applicability to multiple elite maize inbreds. The use of developmentally regulated promoters driving *Cre* enabled auto-excision of morphogenic genes, but resulted in low transformation frequency and QE recovery. These limitations were addressed using heat-shock inducible promoters driving expression of Cre, that resulted in higher frequencies of T0 transformation, gene-excision and QE recovery.

### Excision via developmentally-regulated promoters

The presence of morphogenic genes in transgenic events is undesirable because of unpredictable phenotypes (Lowe et al., 2016). Auto-excision of morphogenic genes occurs early in the transformation process which enables trait evaluation in T0 generation and reduces attrition due to T0 sterility. We evaluated several auto-excision designs, using *Cre* driven by various promoters. These included seven different developmentally regulated (embryo or meristem) promoters, the constitutive maize ubiquitin (*ZM-Ubi*) promoter, and the *Agrobacterium* nopaline synthase (*Nos*) promoter (Table 1). To facilitate excision, the morphogenic genes (*Wus2* and *Bbm*) and the *Cre* gene cassette were flanked with a single pair of directly oriented loxP sites (Figure 1 A). The resulting excised events following auto excision is depicted in Figure 1B. We evaluated two different inbreds (HC69 and PH2RT) to identify *pro:Cre* combinations that produced high frequencies of both transformation and excision. Molecular event data is presented in Table 2. All constructs tested produced stable transgenic events with some number of properly excised events. The *Ole_pro_:Cre* had the highest transformation frequencies (27.2%-37.1%), while the *Glb1_pro_:Cre* construct produced events with higher QE frequencies (8.6%-18.4%).

**Figure 1.**
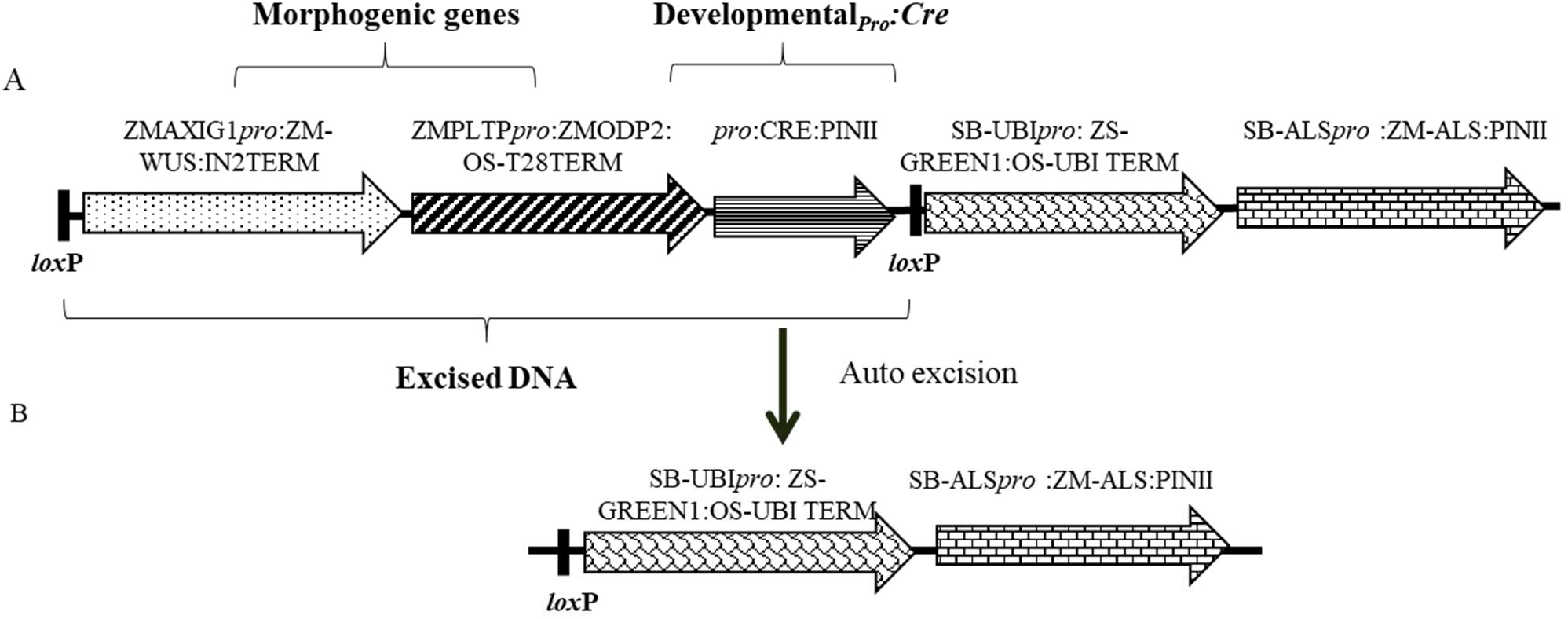
Schematic representation of an auto-excision construct design used for testing different developmentally regulated or stress-inducible promoters to achieve excision of morphogenic genes. A) The excision construct with different promoter combinations driving *Cre* expression (represented by *pro:CRE*) and the DNA fragment to be excised flanked by two directly oriented loxP recombination sites. B) The excised product following auto-excision. Refer to Table S-1 for description of construct components used in T-DNA construction.

**Table 1.** List of the promoters, their source, and their expression pattern in plants.

| Promoters | Source | Expression | Reference |
| --- | --- | --- | --- |
| *Kn1* | Maize | Apical Meristem | Gen bank AY312169 |
| *Lec1* | Maize | Early Embryo | (Shane, 2007) |
| *End2* | Maize | Early Embryo | (Casper et al., 2005) |
| *Ltp2* | Maize | Early Embryo | (Kalla et al., 1994) |
| *Glb1* | Maize | Late Embryo | (Liu et al., 1998) |
| *Ole* | Maize | Late Embryo | (Anand et al., 2017b) |
| *Rab17* | Maize | Late Embryo/Stress | (Busk et al., 1997) |
| *Nos* | *Agrobacterium tumefaciens* | Constitutive | (An, 1986) |
| *Ubi<sub>pro</sub>* | Maize | Constitutive | (Christensen et al., 1992) |
| *Hsp17.7* | Maize | Heat shock inducible | (Anand et al., 2017a) |
| *Hsp26* | Maize | Heat shock inducible | (Anand et al., 2017a) |
| *Rab21* | *Seteria itallica* | Drought inducible | Previously unpublished Corteva Agriscience sequence Si026926m |
| *Drp12* | *Brachypodium distachyon* | Drought inducible | Previously unpublished Corteva Agriscience sequence Bradi3g43870.1 |
| *Drp1* | *Brachypodium distachyon* | Drought inducible | Previously unpublished Corteva Agriscience sequence Bradi1g37410.1 |

**Table 2.**
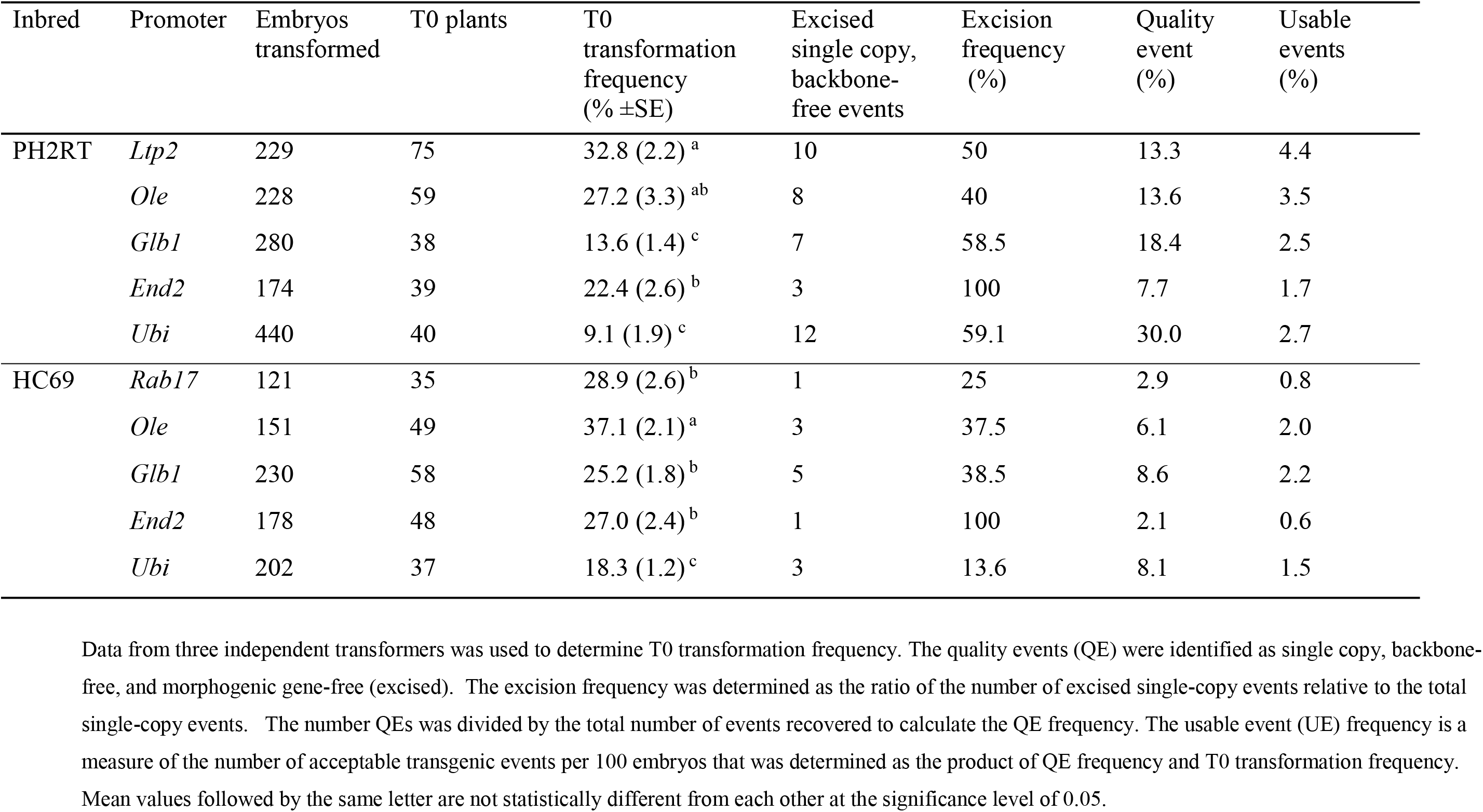
Transformation results with different developmentally regulated promoters driving *Cre* expression for auto-excision of morphogenic genes using construct design described in Figure 1. Data presents the T0 transformation frequency, qPCR detection of the number of excised events and the quality event frequency in two different inbreds, PH2RT and HC69.

### Excision via marker gene activation

Although we achieved auto-excision with all developmentally regulated promoters tested, even for the best construct the usable events rate was around 2% and 80-90% of events were not excised quality events. To improve efficiency, we designed constructs with SMG that was activated only upon excision of the morphogenic genes. This approach selects directly for excised events and was expected to increase QE frequency. A similar construct design was previously used to optimize tissue culture conditions for recovering high quality maize transgenic events (Chu et al., 2019). A schematic design of the construct is depicted in Figure 2A and the quality excised product in Figure 2B. For these experiments, either the *Glb1* or the *Ole* promoters were used to drive *Cre* expression for evaluation of excision-activated marker gene selection. The data from side-by-side testing of these two promoters using the construct design described in Figure 2 are summarized in Table 3. The construct containing *Glb1_pro_:Cre* improved T0 transformation and QE frequencies (1.8 and 1.4-fold), compared to the developmentally regulated gene-excision approach. When *Ole_pro_:Cre* was used, the T0 transformation frequency was similar (>l.l-fold) while the QE frequency increased approximately 1.7-fold. The excision frequency was higher when excision-activated selection was used, with excision frequencies of 53.3% (*Ole_pro_:Cre*) and 68.4% (*Glb_pro_:Cre*) when compared to the previous approach. Additionally, no null events (escapes) were identified by qPCR analysis.

**Figure 2.**
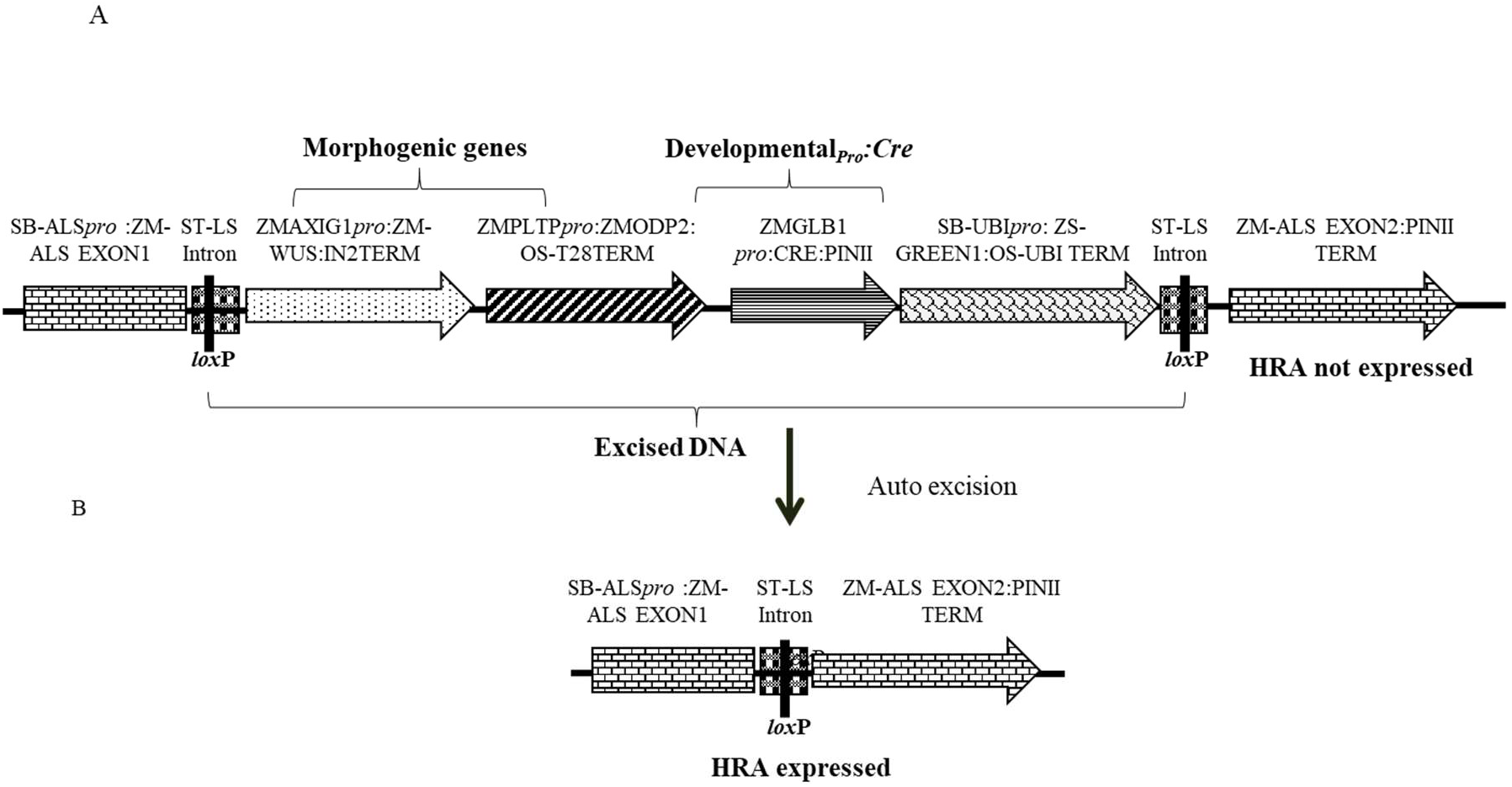
Schematic representation of an auto-excision construct design used for testing developmentally regulated promoters driving *Cre* expression (represented by *pro:CRE*) for excision-activated SMG expression. A) An excision-activated selectable marker construct design with the DNA fragment to be excised flanked by two directly oriented loxP recombination sites. B) Following excision, the *HRA* gene is activated and events are selected on a media supplemented with 0.1 mg/L imazapyr. Refer to Table S-1 for description of construct components used in T-DNA construction.

**Table 3.** Transformation results from excision-activated marker gene selection using either the *Glb1_pro_* or the *Ole_pro_* driving *Cre* expression using construct design described in Figure 2. Data presents the T0 transformation frequency, qPCR detection of the number of excised events and the quality event frequency in maize inbred HC69.

| Promoter | Embryos transformed | T0 plants | T0 transformation frequency (% ±SE) | Total single copy events | Excised single copy, backbone-free events | Excision frequency (%) | Quality event (%) | Usable events (%) |
| --- | --- | --- | --- | --- | --- | --- | --- | --- |
| *Glb1* | 126 | 57 | 44.7 (2.8)<sup>a</sup> | 19 | 13 | 68.4 | 13.3 | 5.6 |
| *Ole* | 112 | 45 | 40.2 (1.9)<sup>a</sup> | 15 | 8 | 53.3 | 8.8 | 3.6 |

5 Data from two independent transformers was used to determine T0 transformation frequency. Quality events (QE) were identified as single copy, backbone-free, 6 and morphogenic gene-free (excised). The excision frequency was determined as the ratio of the number of excised single-copy events relative to the total single- 7 copy events. The number QEs was divided by the total number of events recovered to calculate the QE frequency. The usable event (UE) frequency is a 8 measure of the number of acceptable transgenic events per 100 embryos that was determined as the product of QE frequency and transformation frequency. 9 Mean values followed by the same letter are not statistically different from each other at the significance level of 0.05.

The *Glbpro:Cre* construct design was further evaluated in two additional inbreds, PH84Z and PH85E, alongside HC69 for comparison (Table 4). QEs were recovered in all three inbreds, which were free of the morphogenic genes with no escapes. Excision frequency was similar (55%-61%) across all the inbreds; QE frequencies varied by genotype: 8.7% (HC69), 27.7% (PH85E) and 6.7% (PH84Z) leading to differences in usable quality event frequency (UE, quality events per 100 embryos): 4.3% (HC69), 3.6% (PH85E) and 1.9% (PH84Z).

**Table 4.**
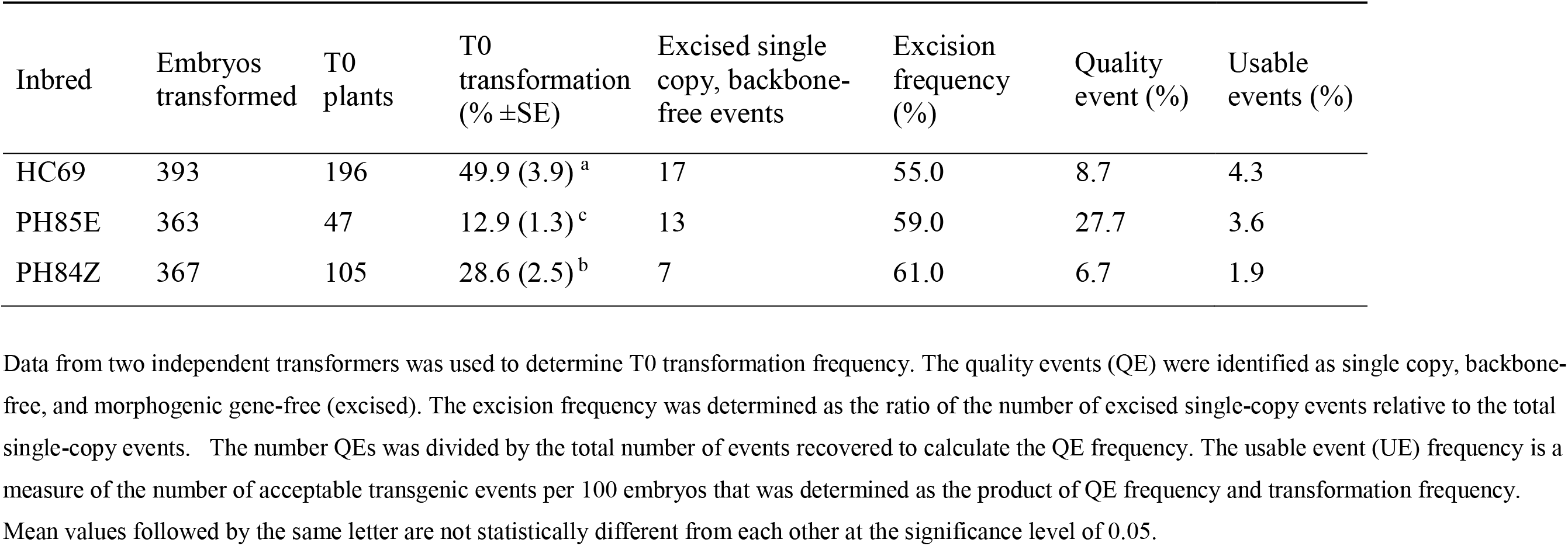
Transformation results from excision-activated marker gene selection using *Glbpro* driving *Cre* expression using construct design described in Figure 2. Data presents the T0 transformation frequency, qPCR detection of the number of excised events and the quality event frequency in three maize inbreds (HC69, PH85E, and PH84Z).

**Table 5.** Transformation results from screening of six different inducible promoters driving *Cre* expression for controlled gene excision. For this study, three different conditions were evaluated: two heat shock treatments (37°C for 1 day and 42°C, 2h/day for 3 consecutive days) and no heat (control). Data presents the qPCR detection of the number of excised events and excision frequency across the different promoters, and a control construct without the *Cre* gene, in maize inbred HC69.

| Promoter | Control |  |  |  | 37°C, 1 day |  |  |  | 42°C, 2h/day for 3 days |  |  |  |
| --- | --- | --- | --- | --- | --- | --- | --- | --- | --- | --- | --- | --- |
|  | T0 |  | Excision |  | T0 |  | Excision |  | T0 |  | Excision |  |
|  | Embryos | plants | QE | (%) | Embryos | plants | QE | (%) | Embryos | plants | QE | (%) |
| *Hsp17.7* | 455 | 59 | 5 | 27.8 | 50 | 6 | 2 | 66.7 | 50 | 20 | 4 | 100 |
| *Hsp26* | 450 | 98 | 0 | 0.0 | 50 | 5 | 0 | 0 | 50 | 21 | 3 | 43 |
| *Rab17* | 455 | 127 | 1 | 3.4 | 50 | 10 | 0 | 0 | 50 | 18 | 0 | 0 |
| *Rab21* | 455 | 101 | 8 | 36.4 | 50 | 13 | 1 | 100 | 50 | 20 | 0 | 0 |
| *Drp12* | 450 | 79 | 2 | 11.1 | 50 | 16 | 0 | 0 | 50 | 22 | 2 | 66.7 |
| *Drp1* | 438 | 90 | 8 | 27.6 | 50 | 8 | 0 | 0 | 50 | 27 | 5 | 45.5 |
| Control (no *Cre*) | 450 | 182 | 0 |  |  |  |  |  |  |  |  |  |

7 Data from two independent transformers was used to determine T0 transformation frequency. The quality events (QE) were identified as single copy, backbone- 8 free, and morphogenic gene-free (excised). The excision frequency was determined as the ratio of the number of excised single-copy events relative to the total 9 single-copy events.

### Excision via stress-inducible promoters

To further improve efficiency, a series of stress-inducible promoters were tested for excision of morphogenic genes. The promoters were selected from a set of genes induced by heat (maize *Hsp17.7* and *Hsp26*) and drought (*ZmRab17, SiRAB21, BdDRP1*, and *BdDRP12*). The construct design is identical to that described in Figure 1, where stress-inducible promoters drive *Cre* expression as represented by *pro:Cre*. The different steps in the transformation process, selection immature embryo infection, In preliminary screening, embryos derived from HC69 were infected with one of the six constructs and, subsequently subjected to one of three different conditions: no heat shock (control), heat shock at 37°C for 1 day, or 42°C for 2h/day for 3 consecutive days. The different steps in maize immature embryo transformation process, included embryo infection with *Agrobacterium* strain continuing the construct (Figure 3A), selection of transgenic events on media supplemented with selectable marker (Figure 3B), heat-shock treatment step (Figure 3C), regeneration of events on media with selection pressure (Figure 3D) and rooting (Figure 3E), before the events were sent to greenhouse. The auto-excision frequencies under induced and non-induced conditions were determined by qPCR analysis. Somatic embryos on maturation media (18 dpi) with 0.1 mg/L imazapyr were subjected to one of the heat conditions and moved onto a rooting media with 0.1 mg/L imazapyr following heat shock (Figure 3D).

**Figure 3.**
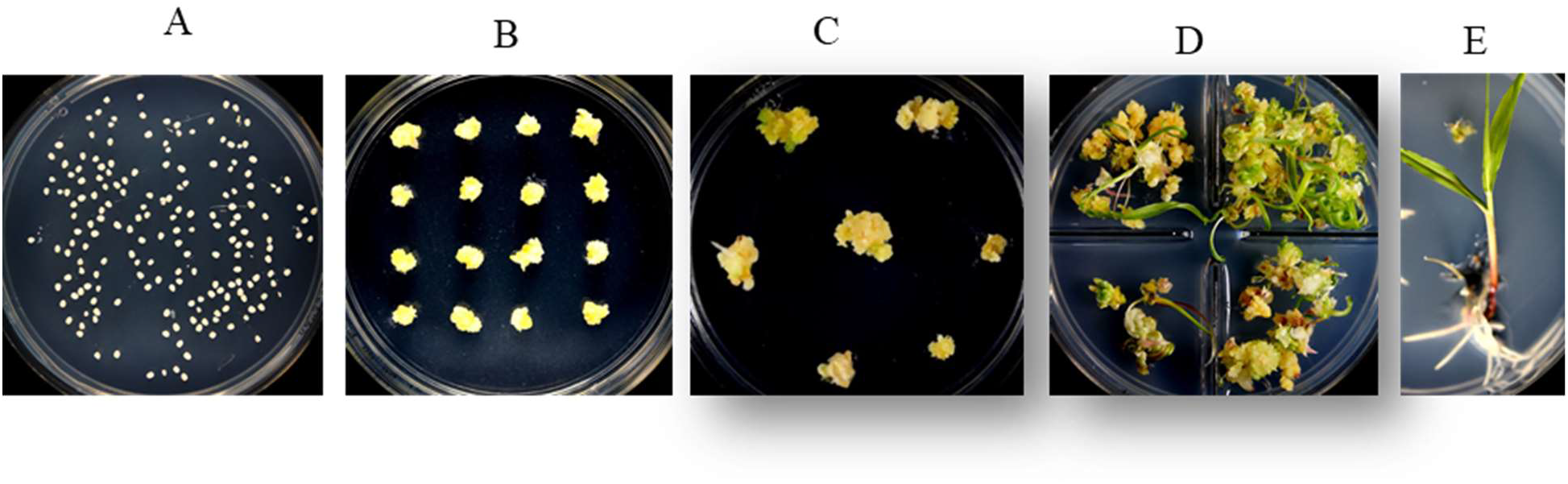
The different stages in rapid maize transformation and heat shock treatment. A) immature zygotic embryos are isolated and infected with *Agrobacterium tumefaciens*, (B) transgenic somatic embryos are placed for 3 weeks on selection media based on selectable marker used (*HRA* or *NPTII*), (C) somatic embryos are heat shock treated and transferred to maturation media, (D) transgenic plants are regenerated without selection pressure for 2 weeks and, (E) regenerated plants are placed on a rooting media for 2-3 weeks.

All promoters except *Hsp26* were leaky under non-induced conditions, resulting in gene-excision rates from 3.4% (*Rab17pro*) to 36% (*BdRab21pro*) compared to zero in the Cre-minus construct. For a subset of the promoters (*Hsp1.7, Hsp26, Drp1* and *Drp12*), higher excision frequencies ranging from 43% to 100%, were observed in the 42°C, 2h/day for 3 days heat treatment. Longer exposure of the somatic embryos at 37°C adversely effected T0 event recovery, compared to a short pulse of heat shock at 42°C (2hr/day for 3 days). Based on the recovery of excised T0 events with *Hsp26_pro_* construct at 42°C treatment compared to 37°C treatment, this promoter appeared to be induced only at higher temperatures.

Additional experiments were performed to further evaluate gene excision and optimize heat shock conditions using three of the inducible promoters (*Hsp17.7, Hsp26* and *Drp12*). HC69 embryos infected with the three constructs were subjected to heat shock treatment at the maturation stage (Figure 3C). One of three different treatments were applied 1) no heat shock (control), 2) 42°C for 2h and 3) 42°C, 2h on 3 consecutive days to determine frequencies of excision and UE recovery (Table 6). Consistent with the previous observation, *Hsp17.7_pro_* driving *Cre* expression under both heat treatments resulted in higher excision rates (62.5%-69.2%) resulting in higher UE rates (10 to 18) compared to *Hsp26_pro_* and *Drp12_pro_*. Based on the data we identified *Hsp17.7_pro_* as the preferred promoter for auto-excision with heat shock of 42°C for 2h.

**Table 6.** Transformation results optimizing the heat shock conditions for controlled gene excision using three inducible promoters driving *Cre* expression. The three different conditions evaluated were: no heat (control) and two heat shock treatments (42°C for 2h and 42°C, 2h/day for 3 consecutive days). The data presents the qPCR detection of the number of excised events and excision frequency across the different promoters in the study as compared to a control construct without the *Cre* gene in maize inbred HC69.

| Promoter | Control |  |  |  |  | 42°C, 2h |  |  |  |  | 42°C, 2h/day for 3 days |  |  |  |  |
| --- | --- | --- | --- | --- | --- | --- | --- | --- | --- | --- | --- | --- | --- | --- | --- |
|  | Embryos | T0 | QE | Excision | UE | Embryos | T0 | QE | Excision | UE | Embryos | T0 | QE | Excision | UE |
|  |  | Plants |  | frequency | % |  | plants |  | frequency | (%) |  | plants |  | frequency | (%) |
|  |  |  |  | (%) |  |  |  |  | (%) |  |  |  |  | (%) |  |
| *Hsp17.7* | 50 | 18 | 1 | 12.5 | 2 | 50 | 17 | 5 | 62.5 | 10 | 50 | 15 | 9 | 69.2 | 18.0 |
| *Hsp26* | 50 | 18 | 0 | 0 | 0 | 50 | 21 | 2 | 42.2 | 4.0 | 50 | 9 | 2 | 66.7 | 4.0 |
| *Drp12* | 50 | 11 | 1 | 25 | 2 | 50 | 9 | 1 | 50.0 | 2.0 | 50 | 14 | 1 | 20.0 | 2.0 |

Data from two independent transformers was used to determine T0 transformation frequency. The quality events (QE) were identified as single copy, backbone-free, and morphogenic gene-free (excised). The excision frequency was determined as the ratio of the number of excised single-copy events relative to the total single-copy events. The number QEs was divided by the total number of events recovered to calculate the QE frequency. The usable event (UE) frequency is a measure of the number of acceptable transgenic events per 100 embryos that was determined as the product of QE frequency and transformation frequency. Mean values followed by the same letter are not statistically different from each other at the significance level of 0.05

### Optimization of heat-shock conditions to improve auto-excision

Further experiments were designed with *Hsp17.7_pro_* and *Hsp26_pro_* to optimize excision conditions. After three weeks of selection, somatic embryos at the maturation stage (Figure 3) were subjected to one of three different heat conditions 1) 42°C, 2h/day for 2 d, 2) 42°C for 24h, or 3) 45°C for 2h/day to determine frequencies of excision and UE. Across the treatments, transformation frequencies ranged from 35%-54.9%, except in the 42°C for 24h treatment of embryos with *Hsp17.7_pro_* driving *Cre* expression, which was lower (Table 7). The heat treatments increased excision rates, which varied with the conditions applied. Of the two *Hsp* promoters tested, *Hsp17.7_pro_* resulted in events with higher excision frequency (75% at 42°C for 24h and 76.6% at 45°C for 2h) compared to *Hsp26_pro_* (66.7% and 61.9%). The treatment, 45°C for 2h worked best for both *Hsp* promoters.

**Table 7.** Optimizing heat shock conditions for controlled gene excision using heat shock promoters *Hsp17.7* and *Hsp26* driving *Cre* expression as shown in Figure 5. Four different conditions were evaluated side-by-side using split ears including no heat (control) and three heat shock treatments (42°C, 2h/d for 2d; 42°C/24h; and 45°C/2h). Transformation results and qPCR detection of the number of excised quality events, frequencies of excision and usable event are presented.

| Promoter | Treatments | Embryos transformed | T0 plants | T0 transformation (% ±SE) | Quality events | Excision frequency (%) | Usable event (%) |
| --- | --- | --- | --- | --- | --- | --- | --- |
| *Hsp17.7* | none | 102 | 56 | 54.9 (4.4)<sup>a</sup> | 6 | 33.3 | 5.9 |
|  | 42°C, 2h/d, 2d | 102 | 39 | 38.2 (2.1)<sup>b</sup> | 9 | 56.3 | 8.8 |
|  | 42°C/24h | 102 | 16 | 15.7 (1.8)<sup>c</sup> | 6 | 75.0 | 5.9 |
|  | 45°C/2h | 102 | 50 | 49.0 (3.2)<sup>a</sup> | 14 | 76.6 | 13.7 |
| *Hsp26* | none | 100 | 53 | 53.0 (4.0)<sup>a</sup> | 1 | 5.6 | 1.0 |
|  | 42°C, 2h/d, 2d | 100 | 35 | 35.0 (1.2)<sup>b</sup> | 12 | 66.7 | 12.0 |
|  | 42°C/ 24h | 100 | 41 | 41.0 (2.2)<sup>b</sup> | 10 | 66.7 | 10.0 |
|  | 45°C/2h | 100 | 50 | 50.0 (2.3)<sup>a</sup> | 13 | 61.9 | 13.0 |

Data from two independent transformers was used to determine T0 transformation frequency. The quality events (QE) were identified as single copy, backbone-free, and morphogenic gene-free (excised). The excision frequency was determined as the ratio of the number of excised single-copy events relative to the total single-copy events. The number QEs was divided by the total number of events recovered to calculate the QE frequency. The usable event (UE) frequency is a measure of the number of acceptable transgenic events per 100 embryos that was determined as the product of QE frequency and transformation frequency. Mean values followed by the same letter are not statistically different from each other at the significance level of 0.05.

### Concurrent elimination of morphogenic and plant selectable marker genes

Next, we developed a strategy that simultaneously excised both the morphogenic genes and the SMG. Two different SMGs, *HRA* and *NPTII* were tested. The construct design was slightly changed to enable excision of the SMG by including it as part of the excised DNA (morphogenic genes and *Cre*) flanked with a single pair of directly oriented loxP sites (Figure 4A) and the resulting excised events are free of SMG (Figure 4B). The binary construct designs with different selectable marker, morphogenic gene and a reporter gene *Zs-GREEN* is illustrated in Figure 4A. Following transformation and selection (either 0. 1 mg/L imazapyr for the *HRA* gene or 150mg/L G418 for the *NPTII* gene), the somatic embryos were heat-shock treated at 45°C for 2h. Transformation data are presented in Table 8. Both *HRA* and *NPTII* constructs produced T0 plants free of morphogenic genes and SMG in the three inbreds tested. With the *HRA* construct, lower frequencies of QEs and UEs were observed and 2-fold more null events were produced compared to the *NPTII* construct. The excision frequency was comparable in both *HRA* and *NPTII* constructs. Irrespective of the differences, both selectable markers produced high frequencies of single copy, backbone-free events which are free of the morphogenic and marker genes.

**Figure 4.**
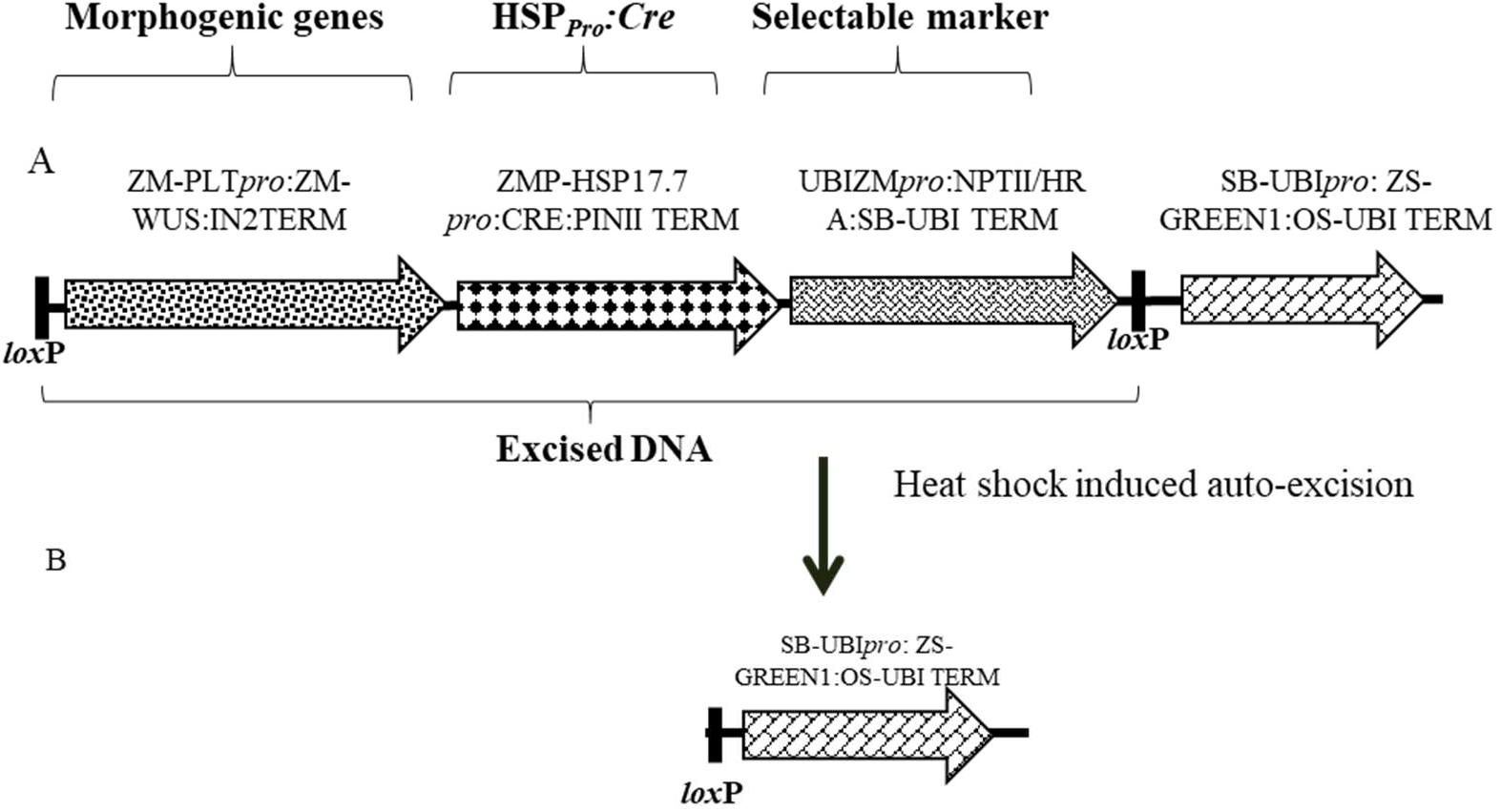
Schematic representation of an auto-excision construct design used for testing elimination of a morphogenic gene and a marker gene using heat shock promoter driving *Cre* expression for controlled gene excision. A) Construct design depicting the order of cassettes including morphogenic genes, *Hsp17.7_pro_* driving *Cre* expression, and the selectable marker (*HRA* or *NPTII*) flanked by directly oriented loxP sites (a) which will be excised upon *Cre* expression. B) Following excision, the DNA piece containing the ZS-GREEN expression cassette is left in the T0 event for visual confirmation of excision. Refer to Table S-1 for description of construct components used in T-DNA construction.

**Table 8.** Transformation results and molecular event data using the *Hsp17.7* heat shock promoter for controlled excision of both morphogenic gene and marker gene in three maize inbreds (HC69, PH85E, and PH84Z). Two different SMGs were evaluated, *HRA* (resistance to the sulfonylurea herbicide ethametsulfuron) and *NPTII* (resistance to antibiotic G418), using the same construct design with the same set of morphogenic genes as shown in Figure 5. Transformation results and qPCR detection of the number of excised quality events, frequencies of excision and usable event are presented.

| Inbred | Selectable marker | Embryos transformed (number) | T0 plants (number) | T0 transformation (%) | Excised single copy, backbone-free events (number) | Excised single copy, backbone-free events (%) | Excision frequency (%) | Usable event (%) | Null (%) |
| --- | --- | --- | --- | --- | --- | --- | --- | --- | --- |
| HC69 | *NPTII* | 315 | 200 | 63.5 | 46 | 23.0 | 87.1 | 14.6 | 17.1 |
|  | *HRA* | 407 | 281 | 69.0 | 45 | 16.0 | 82.3 | 11.1 | 37.3 |
| PH85E | *NPTII* | 219 | 64 | 29.2 | 23 | 35.9 | 96.7 | 10.5 | 15.3 |
|  | *HRA* | 320 | 124 | 38.8 | 31 | 25.0 | 97.2 | 9.7 | 42.5 |
| PH84Z | *NPTII* | 356 | 145 | 40.7 | 19 | 13.1 | 50.4 | 5.3 | 14.2 |
|  | *HRA* | 365 | 169 | 46.3 | 14 | 8.3 | 59.9 | 3.8 | 41.8 |

Data from two independent transformers was used to determine T0 transformation frequency. The quality events (QE) were identified as single copy, backbone-free, morphogenic and marker gene-free (excised). The number of QEs was divided by the total number of events analyzed to calculate the QE frequency. The excision frequency was determined as the ratio of the number of excised single-copy events relative to the total single-copy events. The usable event (UE) frequency is a measure of the number of acceptable transgenic events per 100 embryos that was determined as the product of QE frequency and transformation frequency.

### Progeny analysis

To study the inheritance and segregation of the morphogenic and SMG-free events, we screened single-copy T0 plants free of morphogenic gene and SMG produced from the NPTII construct. Thirteen T0 QE plants, six plants from HC69 and seven plants from PHR84Z, were selected for progeny analysis. These plants were selected and self-pollinated in the greenhouse to enable segregation analysis. Plants from all 13 events produced seeds, 100 to 200 seeds per plant. T1 plants were evaluated for zygosity using qPCR to evaluate copy number of *Cre* and *NPTII* genes (excised DNA). Twelve of the 13 events showed the expected Mendelian inheritance of a single copy T-DNA integration (1:2:1; chi-square p-value>0.05) in the T1 generation (Table 9).

**Table 9.** Observed and expected number of homozygous, hemizygous and null plants for T-DNA integration copy number in in T1 generation of 13 SC excised quality events across two maize inbreds (PH84Z and HC69).

| Inbred | Event ID | Total Plants | Homozygous | Hemizygous | Null | Chi-square | P-value* |
| --- | --- | --- | --- | --- | --- | --- | --- |
| PH84Z | ZMYF66.001.83A | 23 | 7 | 11 | 5 | 0.39 | 0.82 |
|  | ZMCJK9.001.74A | 31 | 10 | 13 | 8 | 0.76 | 0.68 |
|  | ZMCJK9.001.13A | 30 | 8 | 18 | 4 | 2.03 | 0.36 |
|  | ZMCJK9.001.96A | 32 | 6 | 17 | 9 | 0.69 | 0.71 |
|  | ZMCJK9.001.34A | 30 | 10 | 12 | 8 | 1.5 | 0.47 |
|  | ZMCJK9.001.77A | 24 | 5 | 10 | 9 | 2 | 0.36 |
|  | ZMCJK9.001.3A | 27 | 4 | 17 | 6 | 2.07 | 0.35 |
| HC69 | ZMNW4W.001.72A | 23 | 11 | 7 | 5 | 6.65 | 0.03 |
|  | ZMNW4W.001.30A | 31 | 11 | 13 | 7 | 1.83 | 0.39 |
|  | ZMNW32.001.49A | 32 | 4 | 17 | 11 | 3.19 | 0.2 |
|  | ZMNW32.001.58A | 31 | 8 | 14 | 9 | 0.35 | 0.84 |
|  | ZMNW32.001.43A | 32 | 9 | 10 | 13 | 5.5 | 0.06 |
|  | ZMNW32.001.65A | 32 | 9 | 14 | 9 | 0.5 | 0.78 |

- 3 \* No statistically significant deviations identified from expected 1:2:1 (homozygous:hemizygous:null) segregation at 5% level

## DISCUSSION

In maize, direct induction of somatic embryos capable of rapidly germinating from immature embryos (without a callus phase) has been demonstrated using the auxin-inducible promoter *Axig1* driving *Wus2* expression in combination with *Bbm* driven by a maize *PLTP* promoter (Lowe et al., 2018). Continued expression of morphogenic genes results in abnormal phenotypes (Lowe et al., 2016). Therefore, removing morphogenic genes is imperative for accurate construct evaluation and product development and, therefore, a prerequisite for broad application of the technology. Morphogenic gene excision was accomplished using a drought-inducible *Rab17* promoter driving *Cre* recombinase expression (Vilardell et al., 1991). Although this approach was used for successful excision, the requirement for a desiccation step significantly reduced stable event recovery and excision frequency (Lowe et al., 2016).

In order to develop a more efficient system promoters of seven developmentally regulated genes, the *Knotted-1* (*Kn1*) (Bolduc et al., 2012), *Leafy cotyledon1 (Lec?*) (Pelletier et al., 2017), barley *Lipid transfer protein2 (Ltp2*) (Kalla et al., 1994), an early embryo response gene (*End2*) (Casper et al., 2005), *Globulin1* (*Glb1*) (Belanger and Kriz, 1991), and *Olesin* (*Ole*) (Anand et al., 2017b) were evaluated for their ability to express *Cre* and excise morphogenic genes. *Glb1, Ole*, and *End2* promoters unlike inducible promoters did not need either physical or chemical induction for auto-excision. While morphogenic gene removal was observed using developmentally regulated promoters, this generally resulted in lower QE frequencies. A possible explanation is that premature expression caused by early unintended low-level expression from the developmentally regulated promoters resulted in low levels of *Cre* expression.

Developing a method for regenerating events that are free of morphogenic genes using an excision-activated marker gene system may increase excision frequency and QE recovery is described (Chu et al., 2019). In a similar manner, developmentally regulated promoters *Glb1* and *Ole* that are active during late embryo development (Kriz et al., 1990;Anand et al., 2017b), were used to drive *Cre* expression for auto-excision. This strategy resulted in the reconstitution of the *HRA* marker gene, which conferred herbicide resistance (Chu et al., 2019) and would grow in the presence of selective agent. As anticipated, the strategy resulted in improved frequencies of T0 transformation and QE that resulted in approximately a 2-fold increase in UE production. Despite excision of the morphogenic genes and activation of selectable marker, a large proportion of T0 events were multi-copy and non-excised. One possible explanation is the dosage effect of the *HRA* gene on rapid maize transformation, leading to enrichment of events with stable insertions of more than one copy of the transgene. The other possibility is the restricted activation of the developmental promoters leading to partial/incomplete excision, which does not work in rapid maize transformation for enriching quality events.

To achieve controlled expression of recombinases genes for excision, inducible promoters have been an attractive choice. These promoters predominantly fall into two categories; 1) heat shock- or stress-inducible promoters (Kilby et al., 1995;Cuellar et al., 2006;Zhang et al., 2006;Du et al., 2019) and, 2) chemical inducible promoters (Gatz, 1996;Zuo and Chua, 2000). Expressing the recombinase under the control of promoters requiring inducers (heat, osmotic, or chemical) has allowed tighter control of gene expression, while minimizing the negative effect of ectopic gene expression. Among the stress-inducible promoters tested, *Hsp17.7_pro_* and *Hsp26_pro_* produced the best results for auto-excision based on a higher frequency of T0 transformation, gene excision and UE rate. In maize, the regulation of *Hsp* promoters in response to stresses has been described (Pegoraro et al., 2011), including accumulation of *Hsp* proteins under temperatures over 32-33°C (Ristic et al., 1991;Vierling, 1991) and enhanced *Hsp70* synthesis under drought and/or heat (Hu et al., 2010). The heat-inducible auto-excision system was previously described using a construct design that involves *Hsp70pro* driving the *Cre* recombinase for elimination of the SMG (egfp) while a second marker gene, expressing the anthocyanin pigmentation (Rsc) gene, was used for event sorting (Du et al., 2019). While successful, the strategy has limited practical application requiring tracking of transgenes in the T1 generation and subsequent segregation, which is resource-and time-intensive.

Taking a methodological approach, a system was developed to obtain morphogenic gene-free events at high frequencies (66%-77% of the total events generated). The overall strategy was to develop an efficient auto-excision system for rapid maize transformation, with the objective of eliminating both morphogenic and marker genes, that is highly efficient to meet the needs of high throughput maize transformation. The method we developed resulted in the elimination of morphogenic and marker genes at the maturation stage of transformation at high frequencies (ranging from 60%-97%) in multiple elite inbreds. This was achieved by optimizing tissue culture conditions, optimization of heat shock treatment and identifying a versatile SMG. The stably transformed plants were normal, produced seeds and showed stable transmission of the integrated T-DNA to the next generation.

## MATERIALS AND METHODS

### Plant Material

Pioneer temperate maize inbreds (R03, PH2RT, PH85E and PH84Z) were used in this study. All plants used for source immature embryos were grown in the greenhouse. One of the inbred lines (R03) is nonproprietary and publicly available. The other three inbred lines described here are proprietary (PH2RT, PH85E and PH84Z). In order to protect Corteva Agriscience proprietary germplasm, such germplasm will not be made available except at the discretion of Corteva Agriscience and then only in accordance with all applicable governmental regulations.

### Donor material and tissue culture

Seeds were germinated and grown in a greenhouse at temperature set-points of 25.5/20.0°C (day/night), and 16-h daylight. After 21 d, seedlings were transplanted into 5.9 L pots containing a soil-less substrate composed of 38% Canadian sphagnum peat, 51% composted bark, 8% perlite, and 3% vermiculite by volume and adjusted with lime to a pH of 6.0. Maize ears from the Pioneer inbred lines HC69, PH2RT, PH84Z and PH85E were collected from the greenhouse (Johnston, Iowa) at 10 to 11 d after pollination, when the immature embryos were 1.5-2.0 mm in length. Ears were sterilized with 20% Clorox (final sodium hypochlorite concentration of 1.65%) for 15 min and rinsed three times with sterile distilled water.

### Culture media used for transformations and plant regeneration

Briefly, maize immature embryos (1.5-2 mm) were harvested and used for *Agrobacterium-* mediated transformation, using the media, selection and regeneration methods described previously (Lowe et al., 2018;Chu et al., 2019;Hoerster et al., 2020). All media recipes are described by (Lowe et al., 2018;Chu et al., 2019;Hoerster et al., 2020). For selection, 0.1 mg/L imazapyr was supplemented to somatic embryo formation medium or 150 mg/L G418 was substituted for imazapyr.

### *Agrobacterium-mediated* transformation

Constructs used in these experiments are illustrated in Figures 1, 2, and 4 and the individual expression components such as promoters, structural genes and terminators are listed in Table S 1. The materials reported in this article contain selectable markers (*HRA* and *NPTII*) and reporter genes (*ZS-Green* and *Zs-Yellow*) are owned by third parties. Authors may not be able to provide materials including third party genetic elements to the requestor because of certain third-party contractual restrictions placed on the author’s institution. In such cases, the requester will be required to obtain such materials directly from the third party. The authors and authors’ institution do not make any express or implied permission(s) to the requester to make, use, sell, offer for sale, or import third party proprietary materials.

All transformations were done using the thymidine auxotrophic *Agrobacterium tumefaciens* strain LBA4404 THY-containing pVIR9 (Anand et al., 2018) at OD550 of 0.5. The conditions for *Agrobacterium* suspension culture preparation following embryo isolation and infection has been previously described (Lowe et al., 2018;Hoerster et al., 2020). Two selectable markers were used in experiments: *HRA* (Green et al., 2009), a sulfonylurea herbicide resistance marker, driven by the sorghum *Als* promoter for selection with 0.1 mg/L imazapyr in culture medium, or the *Ubipro::NPTII* gene for selection with 150 mg/L G418 in culture medium.

### Excision conditions

For the developmentally regulated *pro::Cre* testing, no optimization was required. These experiments were performed on two inbreds, HC69 and PHR2HT. The initial heat shock treatment for excision involved three different conditions: no heat shock (control), heat shock at 37°C for 1 day, or 42°C for 2h/day for 3 consecutive days, were tested. We further optimized the heat shock condition testing three additional heat treatments 1) 42°C, 2h/day for 2 d, 2) 42°C for 24h, or 3) 45°C for 2h/day to identify a treatment that is best and simple for implementation.

### Molecular analyses

All molecular analysis and transgene copy number determination methods were previously described (Wu et al., 2014;Lowe et al., 2016;Hoerster et al., 2020). qPCR data was used to confirm recombinase-mediated excision based on the absence the transgenes flanked by the loxP sites, determine the copy number of structural genes outside the excision DNA, and to screen for the presence of *Agrobacterium* binary construct backbone integration. Genomic DNA samples were extracted from a single piece (200 ng) of fresh leaf tissue from each plant (Truett et al., 2000). Non-transgenic maize inbred lines were used as the negative controls. Quantification was based on detection of amplified gene sequences using gene-specific forward and reverse primers, along with the corresponding gene-specific FAM™ or Vic®-based MGB fluorogenic probes (Applied Biosystems). The 2-ΔΔCT method (Livak and Schmittgen, 2001) was used to estimate copy number. Events which are single copy for all the transgenes and excised was used to calculate the excision frequency. The events which are excised with a single copy (SC) of all the transgenes without vector backbone integration were defined as a quality event (QE). The usable event (UE) frequency was calculated as transformation frequency times QE frequency. Data collected from different experiments were analyzed separately by analysis of variance (ANOVA), with mean separation by LSD (P=0.05) using JMP Pro 12.2.0 Statistical Discovery software package (SAS Institute Inc., Cary, NC).

## CONCLUSION

Despite the recent progress in developing a rapid maize transformation, the presence of morphogenic genes in the transgenic event have shown to result in pleiotropic phenotypes and is not recommended for transgene testing or commercial product development. The first generation of rapid maize transformation method was designed to improve the transformation rates and to extend transformation capabilities to many genotypes. Subsequently, we demonstrated a viable second-generation alternative, using a mixture of an *Agrobacterium* strains, one with non-integrating *Wus2* gene and the other with a combination of structural genes to regenerate transgenic plants free of morphogenic genes. Even though this simplifies vector construction, however, the process still relies on SMG for recovery of stable transgenic events. This study demonstrated a viable third alternative, relying on inducible promoters for auto-excision of both the morphogenic genes and the SMG in the early stages of maize transformation. The stable transformed plants recovered by this method are free of the morphogenic genes and marker genes, a desirable quality for transgene evaluation and in commercial products.

## AUTHOR CONTRIBUTION STATEMENT

A.A., E.W., L.K., W.G-K., T.J., and N.D.A conceived the research idea, A.A., E.W., L.K., and W.G-K. designed constructs and research, and N.W., MA., HG. and R.L conducted maize transformation and optimization; E.W. and A.A, performed data analysis; A.A., W.G-K. T.J., and N.D.C. wrote the manuscript.

## ACKNOWLEDGMENTS

We thank the internal support groups, Super-Vector (SV) team for their support with vector construction and PCR Analysis and Characterization (PAC) team for molecular event quality analysis. Scott Betts with program support, Terry Hu for maize transformation support. Special thanks to Tracy Fisher and Scott Betts for critical reading of the manuscript and Kara Califf for the art work.

## Abbreviations

*Bbm*: Babyboom
*Cre*: CRE recombinase
HSP: Heat-shock promoters
SMG: Selectable marker-free
Pro: promoters
QE: Quality events
UE: Usable event
*Wus*2: Wuschel2

---

**Table S-1.** Construct components used in T-DNA construction.

| Component type | Label | Description | References |
| --- | --- | --- | --- |
| **Promoters** | *Sb-Als<sub>pro</sub>* | The sorghum ALS promoter | SB-ALS promoter and 5'UTR, DOE-JGI Sbi v3.1, SBChr04, bases 49239164-49240031. DOE-JGI Sbi v3.1 corresponds to Sorghum bicolor BTx623 assembly v3.0.1 and gene annotation v3.1 available from phytozome (http://phytozome.jgi.doe.gov/) . Chromosome 4 of Sbi v3.1 is registered as NCBI accessions NC_012873.2 and CM000763.3 |
|  | *Pltp<sub>pro</sub>* | Maize phospholipid transferase promoter | See GenBank sequence (MN380778) |
|  | *Axig1<sub>pro</sub>* | The maize Axig1 promoter | (Garnaat et al., 2002) |
|  | *Sb-Ubipro* | The sorghum Ubiquitin promoter | (Shane, 2007) |
| **3' Sequences** | *In2-2* | The maize IN2-2 terminator | (Hershey and Stoner, 1991) |
|  | *PINII* | The potato proteinase inhibitor II (pinII) 3' sequence | (An et al., 1989) |
|  | *Os-Ubi 3'* | The rice Ubiquitin terminator | Terminator region of the rice Ubiquitin (Os06g46770.1), unpublished |
|  | *Sb-Ubi 3'* | The sorghum Ubiquitin terminator | (Shane, 2007) |
|  | *Os-T28 3'* | The T28 3' regulatory sequence from *Oryza sativa* | (Bhyri et al., 2014) |
| **Marker genes** | *NPTII* | Maize codon-optimized Neomycin Phosphotransferase II | Previously unpublished Corteva Agriscience sequence |
|  | *HRA* | The maize ALS double mutant gene conferring herbicide resistance | (Green et al., 2009) |
|  | *Zs-YELLOW* | The Zs-Yellow1 N1 gene encoding a yellow fluorescent protein from *Zoanthus sp* | (Matz et al., 1999) |
| **Maize morphogenic genes** | *Zm-Wus2* | The maize *Wuschel2* (*Wus2*) gene | (Lowe et al., 2007) |
|  | *Zm-Bbm* | The maize *Baby boom* gene (*Bbm*) | (Gordon-Kamm et al., 2005) |
| **Recombinase Expression Cassettes** | *Cre* | A maize-optimized *Cre* recombinase gene (originally from the P1 bacteriophage), with an inserted potato LS1 intron | (Odell et al., 1990) |
| **Recombinase Target Sites** | *loxP* | The recombinase target site for the Cre recombinase from *E. coli* | (Odell et al., 1990) |

